# Stromal NRG1 in luminal breast cancer defines pro-fibrotic and migratory cancer-associated fibroblasts

**DOI:** 10.1101/2020.04.06.026971

**Authors:** Mireia Berdiel-Acer, Ana Maia, Zhivka Hristova, Simone Borgoni, Martina Vetter, Sara Burmester, Corinna Becki, Khalid Abnaof, Ilona Binenbaum, Daniel Bethmann, Aristotelis Chatziioannou, Max Hasmann, Christoph Thomssen, Elisa Espinet, Stefan Wiemann

## Abstract

HER3 is highly expressed in luminal breast cancer subtypes. Its activation by NRG1 promotes activation of AKT and ERK1/2, contributing to tumour progression and therapy resistance. HER3-targeting agents that block this activation, are currently under phase 1/2 clinical studies, and although they have shown favorable tolerability, their activity as a single agent has proven to be limited. Here we show that phosphorylation and activation of HER3 in luminal breast cancer cells occurs in a paracrine manner and is mediated by NRG1 expressed by cancer-associated fibroblasts (CAFs). Moreover, we uncover an autocrine role of NRG1 in CAFs. This occurs independently of HER3 and results in the induction of a strong migratory and pro-fibrotic phenotype, describing a subset of CAFs with elevated expression of NRG1 and an associated transcriptomic profile that determines their functional properties. Finally, we identified Hyaluronan Synthase 2 *(HAS2)*, a targetable molecule strongly correlated with *NRG1*, as an attractive player supporting NRG1 - autocrine signaling in CAFs.

## Introduction

Breast cancer is the leading cause of cancer-related mortality worldwide in females (Torre, Siegel et al., 2016). It is considered a heterogeneous disease that comprises several molecular subtypes based on gene expression analysis or biomarker expression (Goldhirsch, Wood et al., 2011, Sorlie, 2004). The family of human epidermal growth factor receptor (HER) of tyrosine kinases (TK) has four members, HER1/EGFR, HER2, HER3 and HER4, and eleven ligands (Lemmon & Schlessinger, 2010). Overexpression of HER family members favors cancer development, however, it also renders these tumours suitable targets for efficient anticancer therapies (Hynes & Lane, 2005). For instance, monoclonal antibodies (mAbs) such as trastuzumab and pertuzumab are usually employed in HER2 overexpressing subtypes (Arteaga & Engelman, 2014, Roskoski, 2014).

Among the HER family members, HER3 is emerging as an important component in the luminal subtype of breast cancer, which accounts for about 65-70% of all breast tumours (Cejalvo, Martinez de Duenas et al., 2017). In agreement with the observation that HER3 is required for cell survival in the luminal but not the basal mammary epithelium (Balko, Miller et al., 2012), luminal breast tumours present the highest levels of *HER3* mRNA (Fujiwara, Ibusuki et al., 2014, Morrison, Hutchinson et al., 2013). HER3 lacks or has little intrinsic TK activity and needs to form heterodimers with kinase-proficient receptor TKs to be functional. For HER3-positive solid tumours, several HER3-targeting agents have been undergoing clinical evaluation for the last 10 years and currently thirteen mAbs are in phase 1 or 2 clinical studies. In contrast to HER2 inhibitors, HER3 binding antibodies such as lumretuzumab have shown limited clinical efficacy as single agents, but favorable tolerability (Jacob, James et al., 2018) (Meulendijks, Jacob et al., 2016).

The major activating ligand of HER3 is neuregulin 1 (NRG1). In the presence of NRG1, HER3 heterodimerizes with EGFR, HER2 or HER4. These partner molecules induce HER3 tyrosine phosphorylation, binding of adapter molecules and thereby enabling downstream oncogenic signaling prominently via PI3K/AKT, but also MAPK and JAK/STAT pathways. This ultimately leads to tumour progression (Olayioye, Neve et al., 2000, Yarden & Sliwkowski, 2001).

Several lines of evidence indicate that NRG1 contributes to the development and progression of different tumour types and its expression has been correlated with poor prognosis in breast cancer, head and neck squamous cell carcinoma and pancreatic cancer (Kolb, Kleeff et al., 2007, Montero, Rodriguez-Barrueco et al., 2008, Qian, Jiang et al., 2015, Tsai, Shamon-Taylor et al., 2003). The fact that NRG1 is the main activating ligand of HER3, suggests that tumours with high levels of NRG1 could respond better to anti-HER3 targeted therapies (Ocana, Diez-Gonzalez et al., 2016, Ogier, Colombo et al., 2018, Yun, Koh et al., 2018). Indeed, NRG1-autocrine signaling has been described in a subset of human cancers, such as head and neck and melanoma, to predict sensitivity to HER2/HER3 kinase inhibition (Wilson, Lee et al., 2011, Zhang, Wong et al., 2012). In the case of breast cancer, the relevance of NRG1 ligand in mediating resistance has been previously described (Shee, Yang et al., 2018). However, in comparison to other cancer entities, the expression of NRG1 in breast tumour cells is usually low and the gene is frequently silenced by DNA methylation (Chua, Ito et al., 2009). This suggests that an autocrine signaling is unlikely in breast cancer and rather the activation of HER3 in luminal cancer cells might be dependent on NRG1 expressed by cells in the tumour microenvironment.

The tumour microenvironment is typically composed mainly of cancer-associated fibroblasts (CAFs) acompained by immune cells, vascular cells and extracellular matrix (ECM) (Pietras & Ostman, 2010). CAFs are characterized by the expression of activation markers such as αSMA (alpha smooth muscle actin), FAP (fibroblast activation protein), and FSP1 (fibroblast-specific protein 1) (Orimo & Weinberg, 2007), and are a known source of ECM and soluble factors (e.g. growth and inflammatory factors) which impact tumour growth and progression. The potential of CAFs as therapeutic targets or prognostic biomarkers is still under debate, as CAFs appear to represent a heterogeneous group of cells with diverse and even opposing functions that differentially determine tumour fate (Augsten, 2014, Cortez, Roswall et al., 2014).

Here, we study CAF heterogeneity in luminal breast cancer both at the molecular and functional level. Using primary CAFs derived from tumour tissue of luminal breast cancer patients, we demonstrate how heterogeneous expression of NRG1 in CAFs determines response of cancer cells to therapies blocking the HER3 signaling pathway (Meulendijks, Jacob et al., 2017, Schneeweiss, Park-Simon et al., 2018). Additionally, we uncover an autocrine role of NRG1 in promoting migration and proliferation of CAFs, and identified a *NRG1*-correlating transcriptomic network enriched in motility and fibrosis present in CAFs. Finally, we reveal Hyaluronan Synthase 2 (*HAS2*), a targetable molecule, as a supporting player strongly correlating with *NRG1* expression in primary fibroblasts and patient data.

## Results

### *NRG1* is expressed in the stromal compartment of luminal breast cancer

To verify the expression pattern of *HER3* in different breast cancer subtypes, we used the public METABRIC (Curtis, Shah et al., 2012) and TCGA (Cancer Genome Atlas, 2012) gene expression datasets. In accordance with previous reports (Balko et al., 2012), *HER3* showed consistent higher expression in the luminal subtypes in both datasets (Fig S1A). Conversely, the expression of its main ligand *NRG1* was overall lower with higher levels in basal-like subtypes (Fig 1A).

**Figure 1.**
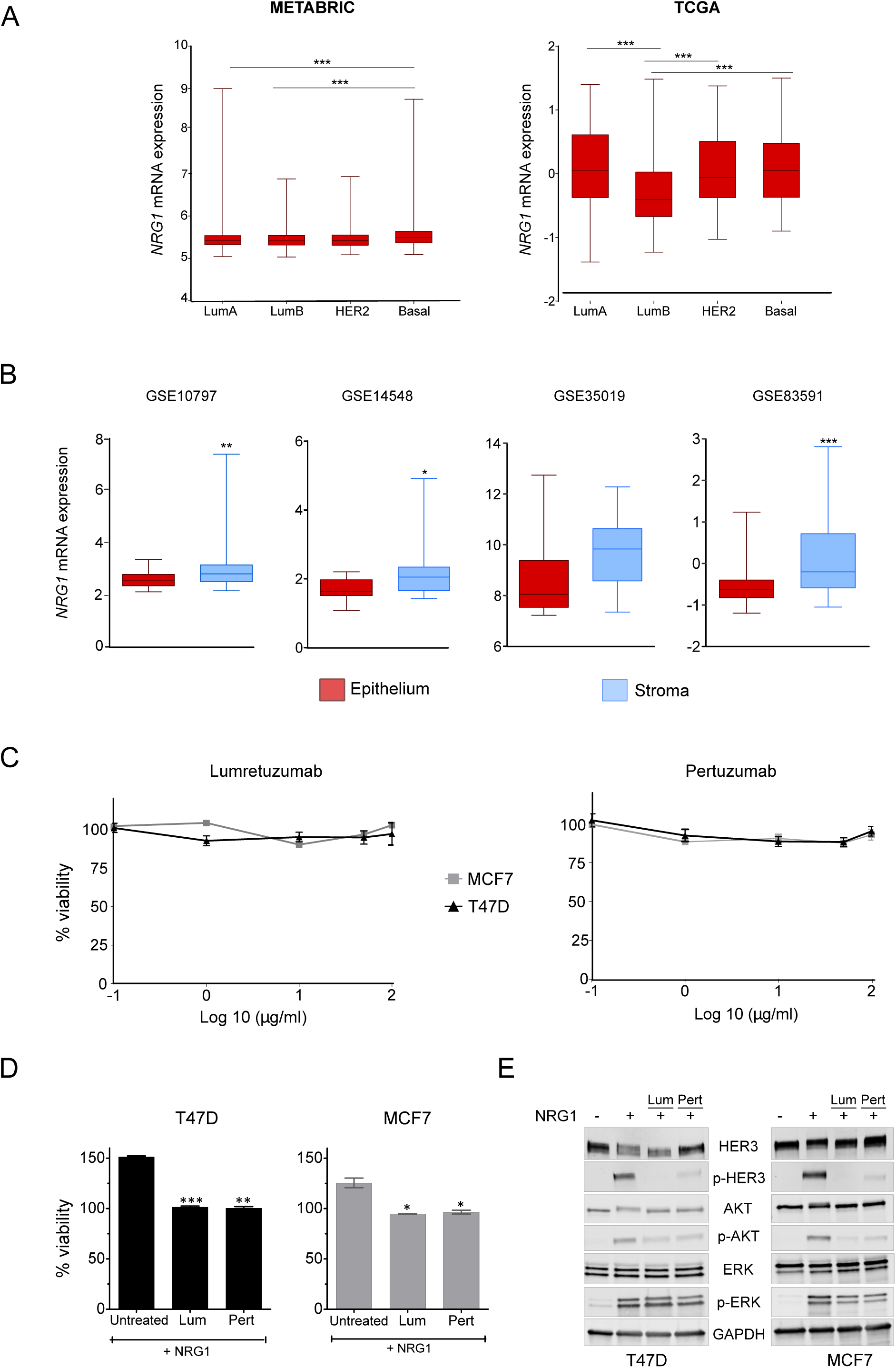
*NRG1* is mostly expressed in the stromal compartment of luminal breast cancer. **A**, expression of *NRG1* in breast cancer subtypes (PAM50) extracted from METABRIC (Curtis et al., 2012) and TCGA (Cancer Genome Atlas, 2012) datasets. Boxes indicate mean +/- quartiles and minimum and maximum values are represented by bars. Only statistically significant comparisons with luminal subtypes are depicted. ANOVA multiple comparison test (\*\*\**P* < 0.001). **B**, expression of *NRG1* in the epithelial and stromal compartment in 4 laser-capture microdissected (LCM) breast cancer datasets (GSE10797, n = 28; GSE14548, n = 14; GSE35019, n = 53; and GSE83591, n = 39). Boxes indicate mean +/- quartiles and minimum and maximum values in bars. Two-tailed paired Student’s t-test (\**P* < 0.05; \*\**P* < 0.01; \*\*\**P* < 0.001). **C**, viability of T47D (black) and MCF7 (grey) measured 72h after treatment with HER3 mAb lumretuzumab or pertuzumab at indicated doses. Values represent median of 3 independent experiments (n = 5). U-Mann Whitney two-tailed test was applied. No significant differences were observed. **D**, ectopic NRG-1β (50ng/mL) was added to T47D and MCF7 cell lines and viability quantified. The untreated control was set to 100% (not shown) and used to normalize the results from conditions where cells had been preincubated for 1 hour with lumretuzumab (Lum) or pertuzumab (Pert) at 10µg/mL. Two-tailed unpaired Student’s t-test (\**P* < 0.05; \*\**P* <0.01, \*\*\**P* <0.001) comparing each treatment with untreated condition. Bars represent average of two independent experiments +/- s.e.m. **E**, representative Western blot showing total levels and phosphorylation of HER3 and downstream effectors AKT and ERK1/2, 5min after addition of NRG-1β (50ng/ml). Some samples were either pre-incubated with mAbs lumretuzumab (Lum) or pertuzumab (Pert) at 10µg/ml for one hour.

Gene expression analysis of bulk tissues comprises mixed signals from different cellular components, masking the contribution of different tumour compartments. Thus, we next explored *NRG1* expression in a collection of breast cancer datasets generated by laser capture microdissection (LCM) of the stromal and epithelial compartments (GSE10797, (Casey, Bond et al., 2009); GSE14548, (Ma, Dahiya et al., 2009); GSE35019 (Vargas, McCart Reed et al., 2012) and GSE83591 (Liu, Dowdle et al., 2017)). In all LCM datasets explored, expression of *NRG1* was higher in the stromal compartment (Fig 1B). This indicates that the stromal cells are the major contributors of *NRG1* expression in breast tumour tissue and suggests that activation of the HER3 pathway in tumour cells preferentially happens in a paracrine manner.

In order to define a proper *in vitro* system for subsequent studies, we analyzed different breast cancer cell lines for expression of the HER family receptors (*EGFR, HER2, HER3* and *HER4).* As in the primary tissue datasets, cancer cell lines from luminal subtypes (T47D, MCF7 and BT474) showed elevated levels of *HER3* (Fig S1B).

We focused on luminal A cell lines T47D and MCF7 to avoid masking of HER3 mediated effects by HER2 overexpression (BT474). To test if luminal A cancer cell lines might be intrinsically addicted to HER3 oncogenic signaling (Weinstein, Begemann et al., 1997), cells were challenged with increasing doses of the therapeutic monoclonal antibody lumretuzumab, that blocks binding of NRG1 to HER3 (Mirschberger, Schiller et al., 2013), or pertuzumab, which blocks HER2/HER3 heterodimer formation (Franklin, Carey et al., 2004). After three days of treatment, viability of cancer cell lines was not affected by HER3 blockage, suggesting no autocrine activation of the HER3 pathway in the luminal A cell lines (Fig 1C). However, HER3 might still be relevant via paracrine activation. To test this, we added ectopic NRG1 to cancer cells that had or had not been pre-incubated with either lumretuzumab or pertuzumab. Whereas the viability of control cells without antibody-incubation was indeed increased by ectopic NRG1, the effect was abolished by pre-treatment with lumretuzumab or pertuzumab (Fig 1D). Additionally, phosphorylation of HER3 and of its main downstream effectors AKT and ERK was efficiently induced by NRG1 in control cells while this was strongly prevented upon pre-treatment with lumretuzumab or pertuzumab (Figs 1E and S1C). Together, these data demonstrate that NRG1 activates HER3 pathway via binding to HER3 in a paracrine manner and that this paracrine activation can be blocked with monoclonal antibodies.

### Primary breast cancer-associated fibroblasts express variable levels of NRG1

Cancer-associated fibroblasts (CAFs) are particularly abundant in the stroma of solid tumours (Walker, 2001). As we had observed that *NRG1* is highly expressed in the stromal compartment of breast tumour tissue (Fig 1B), we next aimed to determine if CAFs are a source of NRG1. To this end, we established primary cultures of CAFs derived from tumour tissues from six breast cancer patients clinically classified as luminal subtype (Table S1).

The isolated cells showed the characteristic fibroblast morphology as well as expression of common CAF activation markers such as αSMA, FAP and fibronectin (Figs 2A-B and S2A-B).

**Figure 2.**
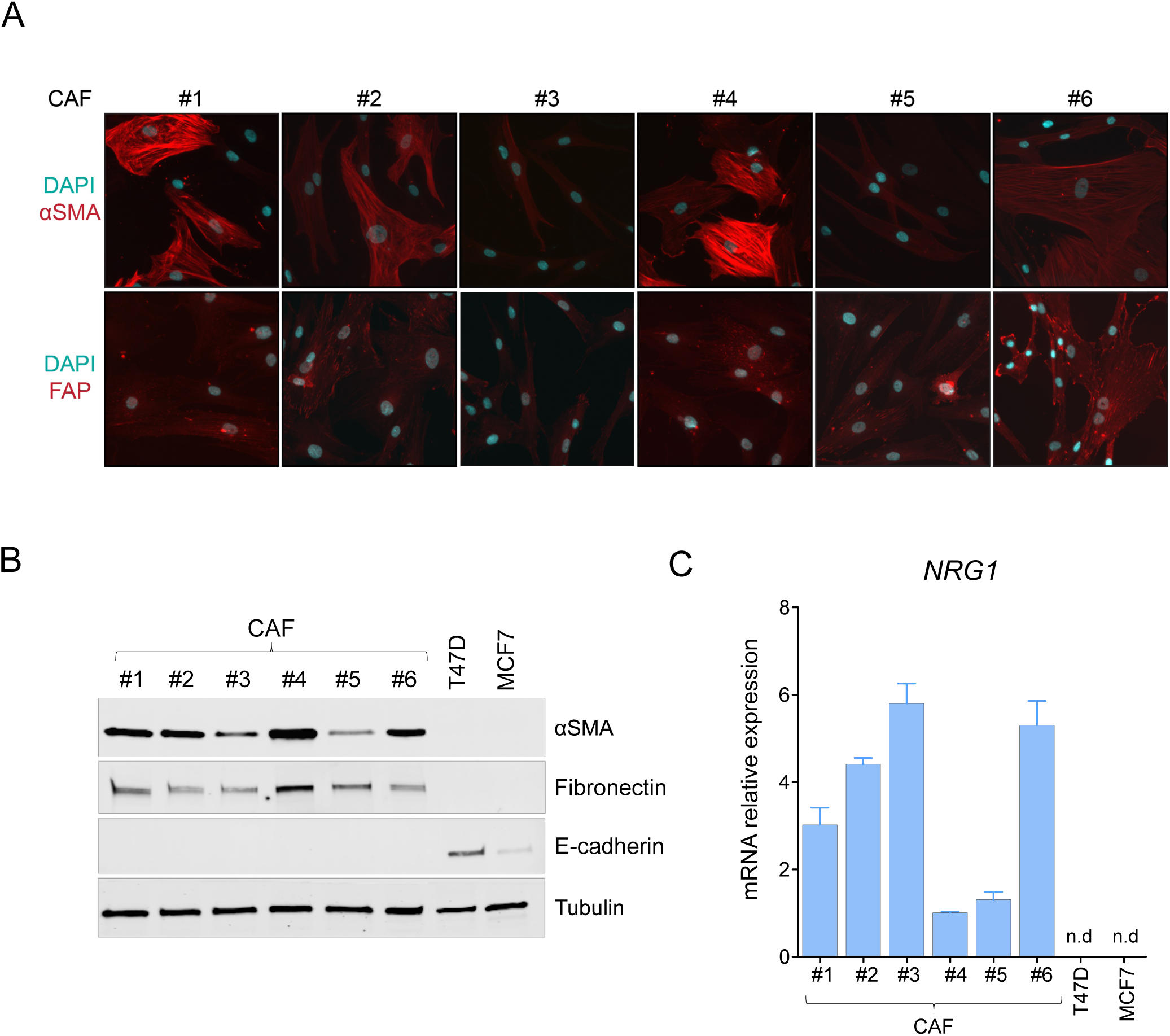
Primary breast cancer-associated fibroblasts express variable levels of *NRG1*. **A**, immunofluorescence of common activation markers αSMA (red, upper panel) and FAP (red, lower panel) in CAFs under study. Nuclei counterstaining with DAPI (blue). Representative images are shown. **B**, Western blot representing protein expression of fibronectin and αSMA in all 6 CAFs under study and in two luminal breast cancer cell lines. The epithelial marker E-cadherin served as marker for epithelial cells. Tubulin was used as a loading control. **C**, relative expression of *NRG1* transcript in all 6 CAFs and in two luminal breast cancer cell lines. Values relative to CAF#4. Bars represent mean +/- s.e.m of two independent experiments.

Next, we analysed mRNA transcript and protein expression levels of *NRG1* in the CAF lines and in the two luminal breast cancer cell lines. In line with the LCM patient data, *NRG1* was expressed by all CAF lines while no expression could be detected in cancer cells (Figs 2C and S2C). Interestingly, NRG1 levels were heterogeneous among the different fibroblasts despite having been isolated from tumours of the same subtype. Collectively, these results demonstrate that NRG1 is expressed by CAFs in the stroma of breast cancer patients and reinforce the concept of a paracrine-driven activation of HER3 in the luminal breast cancer subtype.

### Different levels of NRG1 secreted by CAFs determine activation of HER3 in cancer cells

To ascertain whether different expression of NRG1 by CAFs translates into variable activation of the HER3 pathway in cancer cells, we stimulated T47D and MCF7 cancer cells with conditioned media (CM) from the isolated CAFs, and used lumretuzumab to block ligand-receptor binding.

In order to detect phosphorylation levels of HER3 and its main downstream effectors AKT and ERK in a sensitive and quantitative manner, we applied Reverse Phase Protein Array (RPPA) technology (Sonntag, Schluter et al., 2014). Incubation with ectopic NRG1 as well as blockage with lumretuzumab were used as positive controls (Fig S3A). Phosphorylation of HER3 observed in cancer cells upon stimulation with the different conditioned media was CAF- and cancer cell-dependent, achieving different phosphorylation degrees (Fig 3A, red = maximum, blue = minimum). In both cell lines we observed strong phosphorylation of the HER3 pathway induced by conditioned media from CAF#2 and CAF#3, both of which had higher expression of *NRG1* (Fig 2C). Pre-incubation with lumretuzumab reduced phosphorylation of HER3 and its effectors in all conditions, confirming NRG1-mediated activation of HER3 from the conditioned media.

**Figure 3.**
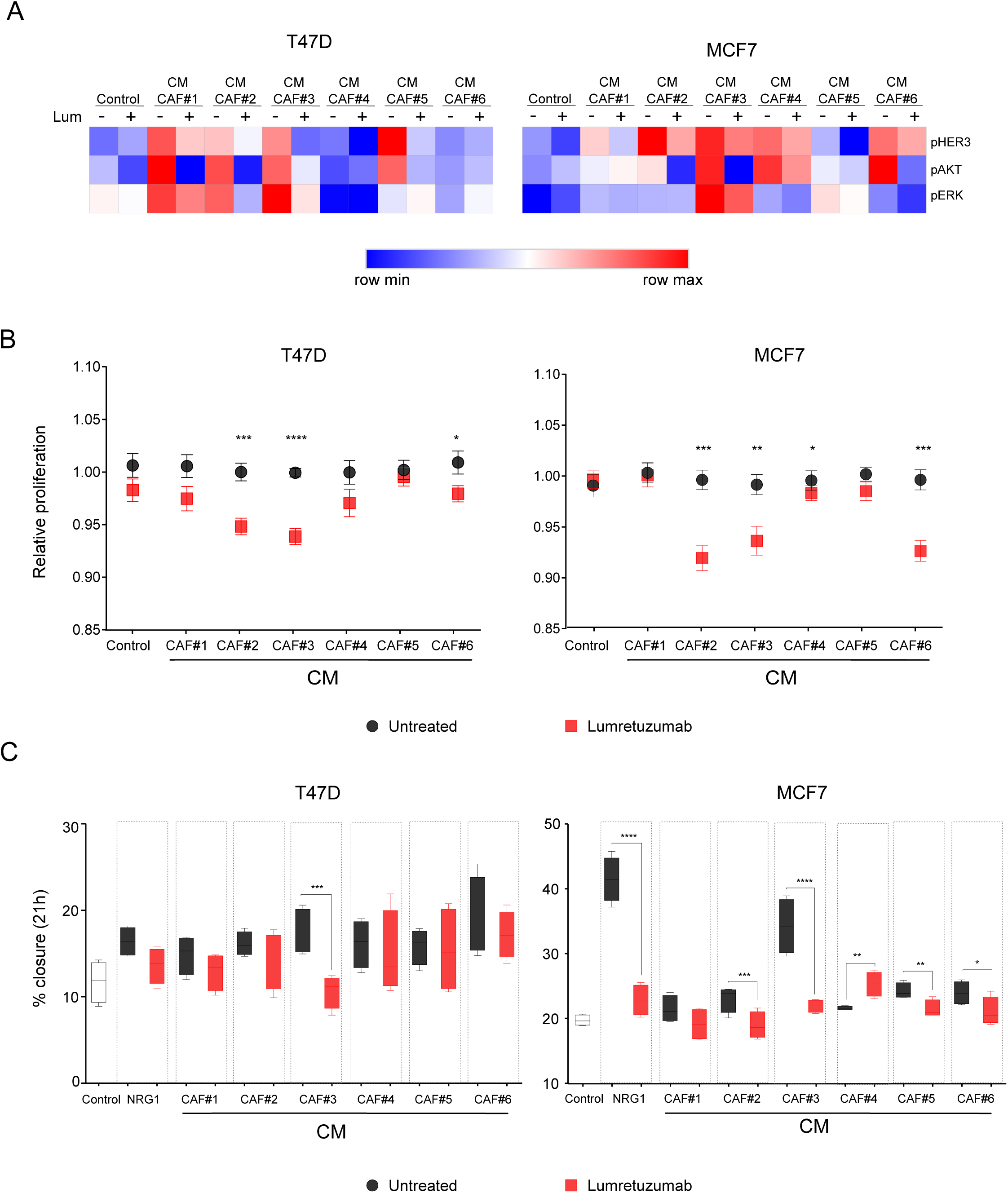
Activation of HER3 in cancer cells by secreted NRG1 is CAF-dependent. **A**, heatmap representing relative phosphorylation values of HER3, AKT and ERK1/2 in T47D and MCF7 cells after addition of CAF conditioned media (CM) for 5min with or without pre-incubation with lumretuzumab (Lum) (10µg/ml). Values represent median of three technical replicates and three biological replicates. Color intensities are ranked per each antibody (red = maximum, blue = minimum). **B**, relative proliferation of T47D and MCF7 cancer cells after 72h with different CAF-CM with lumretuzumab (red squares) or untreated (black circles). Dots represent mean +/- s.e.m of three independent replicates (n = 4). *P* values were determined by two-tailed U-Mann Whitney test for each CM (\**P* < 0.01; \*\**P* < 0.001; \*\*\**P* < 0.0005; \*\*\*\**P* <0.0001). **C**, percentage of closure in a scratch assay of T47D or MCF7 cancer cells, after 21h of treatment with 10μg/ml lumretuzumab (red) or untreated (black), and with conditioned media (CM) of indicated CAFs. DMEM-F12 1% FCS was used as negative control and NRG-1β (50ng/ml) as positive control. Box plots correspond to the mean and s.e.m. of 2 independent experiments (n = 6 technical replicates). Two-tailed U-Mann Whitney test for each CM (\**P* < 0.01; **P < 0.001; \*\*\**P* < 0.0005; \*\*\*\**P* <0.0001).

Next, we tested the ability of NRG1 in the CAF-CM to promote proliferation of cancer cells. The proliferation rate of cancer cells did not follow the same trend as *NRG1* expression (Fig S3B), suggesting alternative proliferation drivers among the different CAFs and/or tumour cells. However, blockage of NRG1-HER3 signaling by lumretuzumab treatment, decreased the proliferation of cancer cells (Fig 3B), specifically when using CM of those CAFs that had promoted higher activation of the HER3 downstream pathways (CAF#2 and #3).

Due to the established role of *NRG1* in epithelial to mesenchymal transition (EMT) and migration processes (Kim, Jeong et al., 2013), we next measured if also migration abilities of the cancer cells were altered in presence of CAF-CM and if these were dependent on NRG1-HER3 activation. Consistently, CAF-CM increased migration of cancer cells and the effect was diminished by treatment with lumretuzumab (Fig 3C). The strongest effect was observed for CAF#3-CM, the CAF culture that expresses the highest level of NRG1. These experiments confirmed that NRG1 from CAFs promotes migration of cancer cells.

Taken together, these results indicate that CAFs isolated from tumour tissue of luminal breast cancer specimens differently activate the HER3 pathway and regulate proliferation and migration of cancer cells via NRG1 secretion/expression.

### Heterogeneous expression of *NRG1* defines two clusters of CAFs

Despite all different cultures of CAFs were derived from luminal breast cancer tissue, they showed variable capacities to activate the HER3 pathway in luminal breast cancer cells via NRG1 (Fig 3A-C). To investigate the possible differences between the isolated CAFs in a global approach, we performed RNA sequencing of the six primary CAF lines. Unsupervised principal component analysis (PCA) revealed that gene expression among fibroblasts was indeed scattering (Fig 4A). To elucidate which genes were contributing to this variance, we analyzed the most significant variable genes (MVG) amongst the different CAF lines. A list of 517 significant genes was defined, with *NRG1* ranked among them (Table S2). Other genes listed as highly heterogeneous were *ACTA2* and *S100A4* coding for αSMA and FSP1 respectively, two well accepted CAF markers, although not correlated with *NRG1* expression (Fig S5B).

**Figure 4.**
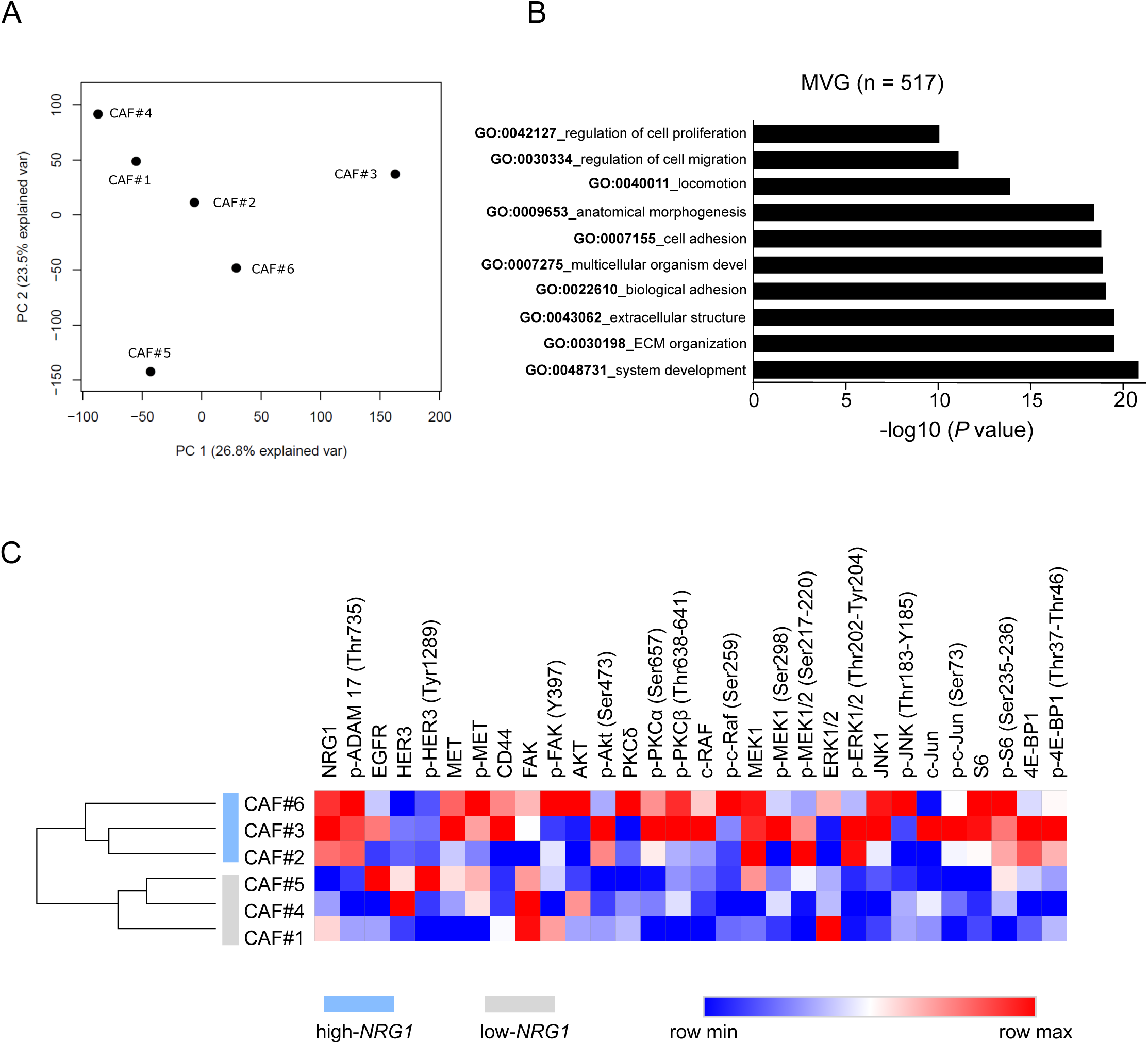
Heterogeneous expression of *NRG1* defines two clusters of CAF. **A**, principal component analysis PC1 and PC2 of CAF lines. **B**, biological processes (Gene Ontology) enriched in CAFs most variable genes (MVG). Graph bars represent adjusted *P* value. **C**, proteomic profile of CAFs based on reverse phase protein array (RPPA). Intensity values (red = maximum, blue = minimum) are ranked per each antibody to compare between samples. Values represent median of three technical replicates.Unsupervised clustering by Euclidian distance separates high and low-NRG1 CAFs (blue and grey respectively).

Next, we performed functional analysis using gene ontology (GO) terms in the Bioinfominer online platform (https://bioinfominer.com) (Koutsandreas T, 2016) for the 517 MVG. Functional categories related to extracellular matrix, cell adhesion and locomotion were significantly enriched (Fig 4B and Table S3). Among the functional categories that were differing in fibroblasts, we found regulation of proliferation. In line with this, nuclei counting of fibroblasts along several days revealed a heterogeneous proliferation degree among the CAFs lines (Fig S4A).

To comprehend to which extent *NRG1* contributed to fibroblasts variability, we split CAFs based on *NRG1* expression into low-*NRG1* (lower than mean: CAF#1, CAF#4, CAF#5) and high-*NRG1* (higher than mean: CAF#2, CAF#3, CAF#6) CAFs (Fig S4B). We perfomed targeted proteomic analysis for the CAFs under study to elucidate if phosphorylation status of effectors of the HER pathway (e.g, ERK1/2, AKT, MET, S6K, ADAM17) were differing among the lines. Strickingly, unsupervised clustering grouped CAFs into two groups demonstrating different activation of the HER3 pathway. Interestingly, NRG1 expression was sufficient to separate those CAFs (Fig 4C). To note, phosphorylation of the transcription factor c-JUN, described as a central molecular mediator in fibrotic conditions (Wernig, Chen et al., 2017) and hyperactivated in high density stroma breast cancer tissue (Lisanti, Tsirigos et al., 2014), was higher in high-*NRG1* CAFs (CAF#2, CAF#3, CAF#6). Moreover, analysis of the transcription factor genes enriched in high-*NRG1* CAFs disclosed c-JUN as the main transcription factor regulating high-*NRG1* transcriptome (Fig S4C and Table S4).

Collectively, these results underline the relevance of NRG1 as a heterogeneous factor discriminating two populations of CAFs in luminal breast tumours.

### *NRG1*-associated transcriptome correlates with migration processes in CAFs

We next wanted to elucidate if different expression of *NRG1* in fibroblasts was associated with specific transcriptional programs. To this end, we performed differential expression analysis between high and low-*NRG1* groups of CAFs. A total of 102 genes were upregulated and 151 were downregulated in the high-*vs* low-*NRG1* CAFs (adj *P* value < 0.05 and absolute logFC > 0.5) (Table S5). Gene Ontology (GO) analysis revealed that genes enriched in the high-*NRG1* CAF group were mainly related with adhesion and motility processes. In contrast, terms enriched in low-*NRG1* CAFs were associated to signaling and metabolic processes (Fig 5A). Based on the list of significant differentially expressed genes (Table 1), we used BioinfoMiner (https://bioinfominer.com) to explore systemic processes and driver genes characteristic of each group. In line with the GO results (Fig 5A), driver genes (*P* value <0,002 and FC> 2) in high-*NRG1* CAFs included *ITGB2, EPHB1* and *HAS2*, known locomotion and extracellular matrix reorganization related genes. In contrast, driver genes in low-*NRG1* CAF included genes such as *PTGIS, TRH, WNT2* or *JAG1*, involved in cell signaling and metabolic processes.

**Table 1:**
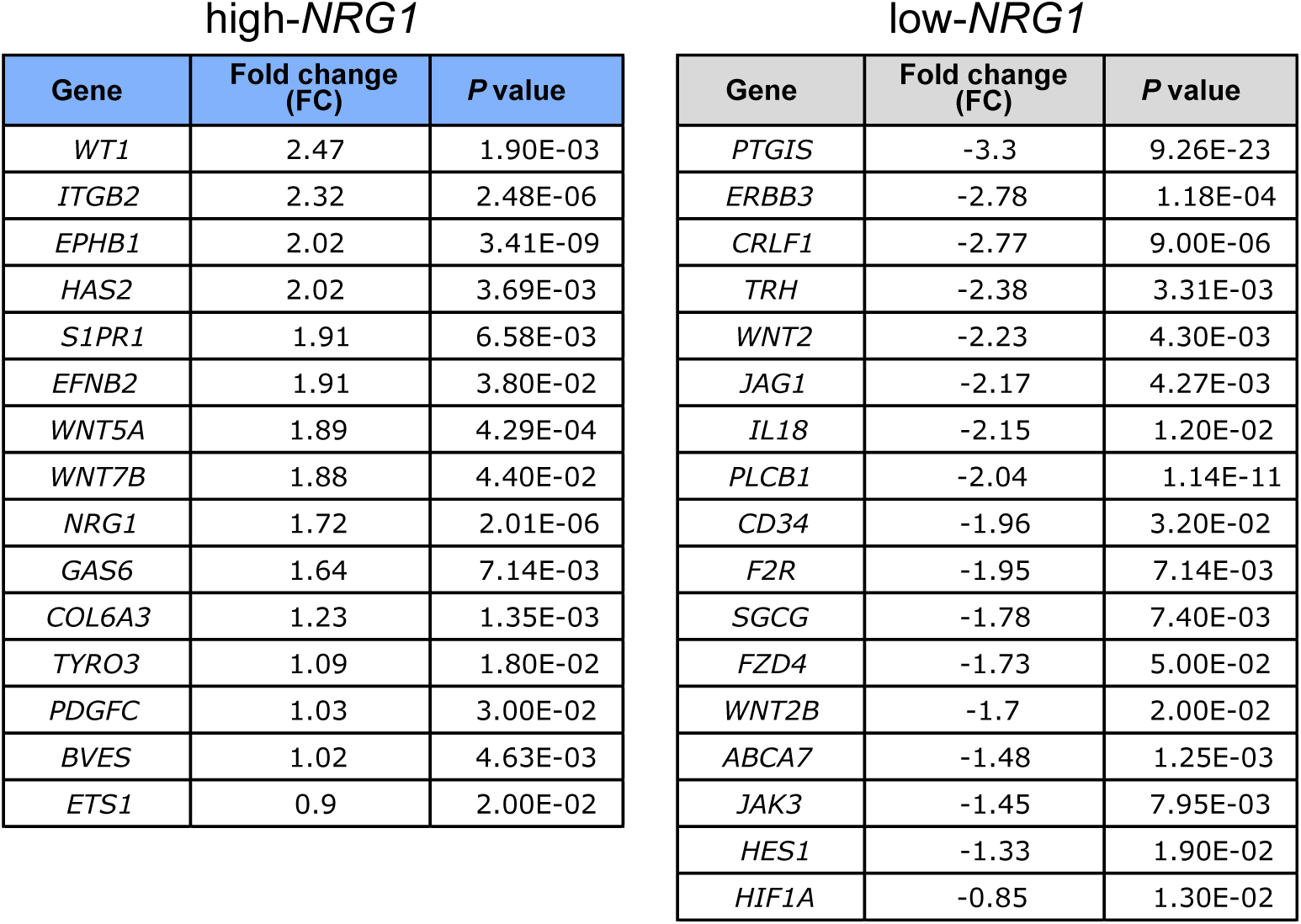
Priorization list of driver genes in each group of CAFs. Analysis of DEG using BioInfoMiner (https://bioinfominer.com) identifed a list of priorization genes as hub nodes for high-*NRG1* and low-*NRG1* CAFs respectively. 15 genes significantly upregulated defining high-*NRG1* CAFs, and 17 genes significantly downregulated in high-*NRG1* CAFs, defining low-*NRG1* CAFs.

| Gene | Fold change (FC) | P value |
| --- | --- | --- |
| *WT1* | 2.47 | 1.90E-03 |
| *ITGB2* | 2.32 | 2.48E-06 |
| *EPHB1* | 2.02 | 3.41E-09 |
| *HAS2* | 2.02 | 3.69E-03 |
| *S1PR1* | 1.91 | 6.58E-03 |
| *EFNB2* | 1.91 | 3.80E-02 |
| *WNT5A* | 1.89 | 4.29E-04 |
| *WNT7B* | 1.88 | 4.40E-02 |
| *NRG1* | 1.72 | 2.01E-06 |
| *GAS6* | 1.64 | 7.14E-03 |
| *COL6A3* | 1.23 | 1.35E-03 |
| *TYRO3* | 1.09 | 1.80E-02 |
| *PDGFC* | 1.03 | 3.00E-02 |
| *BVES* | 1.02 | 4.63E-03 |
| *ETS1* | 0.9 | 2.00E-02 |

low-*NRG1*
| Gene | Fold change (FC) | P value |
| --- | --- | --- |
| *PTGIS* | -3.3 | 9.26E-23 |
| *ERBB3* | -2.78 | 1.18E-04 |
| *CRLF1* | -2.77 | 9.00E-06 |
| *TRH* | -2.38 | 3.31E-03 |
| *WNT2* | -2.23 | 4.30E-03 |
| *JAG1* | -2.17 | 4.27E-03 |
| *IL18* | -2.15 | 1.20E-02 |
| *PLCB1* | -2.04 | 1.14E-11 |
| *CD34* | -1.96 | 3.20E-02 |
| *F2R* | -1.95 | 7.14E-03 |
| *SGCG* | -1.78 | 7.40E-03 |
| *FZD4* | -1.73 | 5.00E-02 |
| *WNT2B* | -1.7 | 2.00E-02 |
| *ABCA7* | -1.48 | 1.25E-03 |
| *JAK3* | -1.45 | 7.95E-03 |
| *HES1* | -1.33 | 1.90E-02 |
| *HIF1A* | -0.85 | 1.30E-02 |

**Figure 5.**
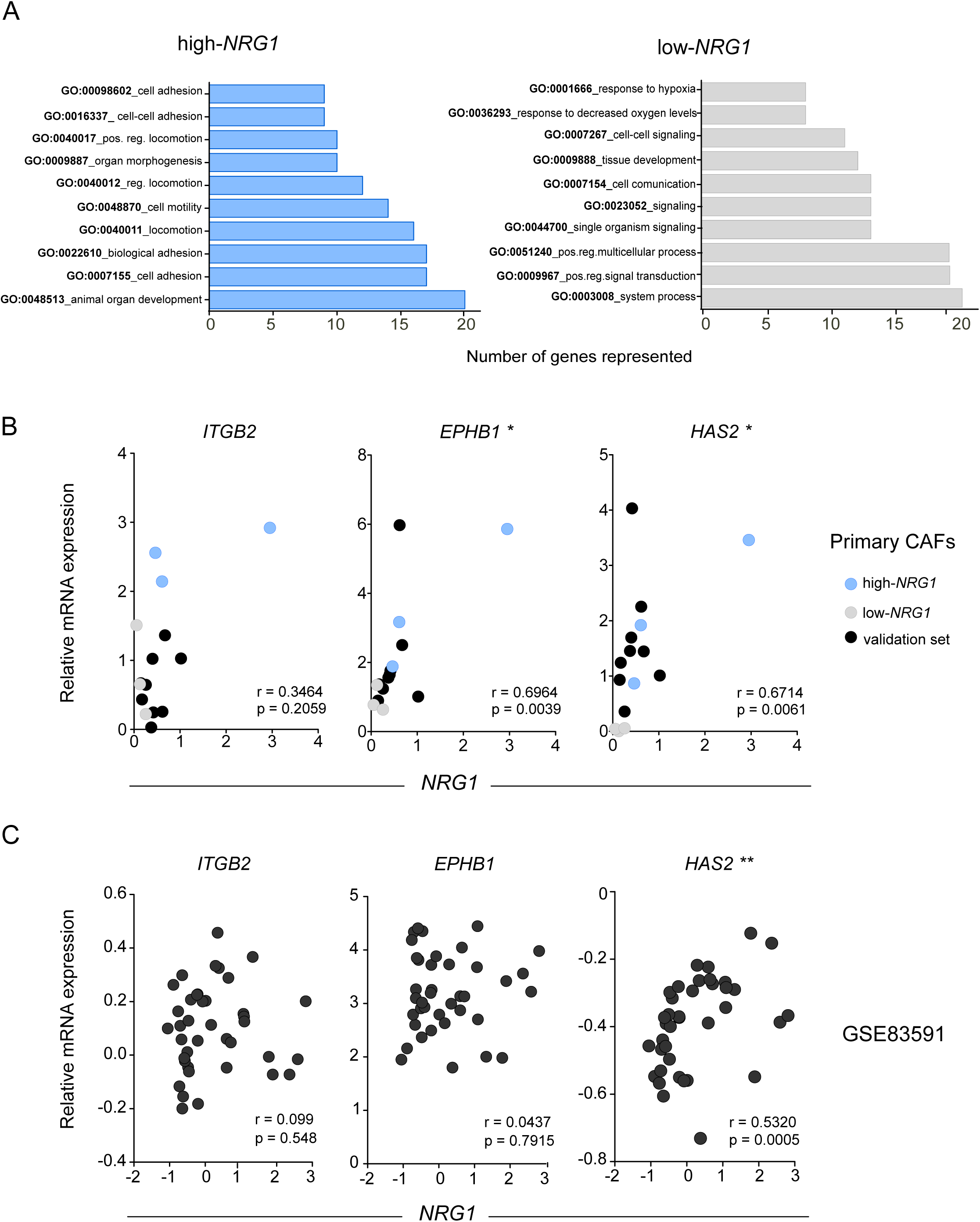
Genes associated to *NRG1* expression. **A**, functional classification of genes by Gene Ontology (GO). Biological processes represented in each group of CAFs. Graph bars showing number of genes with significant *P* value for *NRG1* high and low clusters independently. **B**, expression correlation of *NRG1* and candidate genes quantified by RT-PCR in primary CAFs from two independent sources, six lines from Halle University (discovery set, blue = high-*NRG1* and grey = low-*NRG1*) and another nine from Breast Cancer Now (validation set, black). Each dot represents the average of > 3 independent experiments. Spearman *r* correlation and two-tailed *P*-value are indicated. **C**, expression correlation of *NRG1* and indicated genes in the stroma compartment of the LCM dataset GSE83591 (n = 39). Spearman correlation *r* and *P* value are indicated.

Altoghether, these results determine *NRG1* as a stromal marker discerning two subsets of fibroblasts having different transcriptional programs.

### *HAS2* expression correlates with *NRG1* in the tumour stroma of patient samples

In order to obtain a signature of genes linked with *NRG1* expression in tumour stromal fibroblasts, we selected those genes with the strongest correlation with *NRG1* (Pearson *r* > 0.8), (Fig S5A) and with a FC > 2 in high-*vs* low-*NRG1* CAFs (Table S6).

We validated the correlation of these genes (i.e. *ITGB2, EPHB1* and *HAS2*) with *NRG1* in a second set of primary CAFs from an independent source (https://breastcancernow.org/breast-cancer-research/breast-cancer-now-tissue-bank) (Fig 5B), and we explored their expression in LCM stroma datasets from breast cancer patients (Fig 5C). This analysis confirmed a significant positive correlation between *NRG1* and *HAS2*, which was not present for *ITGB2* and *EPHB1* (Fig 5C). Next, we checked the expression in the TCGA dataset, covering bulk tumour tissue, only considering those samples with tumour purities lower than 50% to select tumours with high stroma content. From the candidates investigated, just *HAS2* was significantly correlated with *NRG1* in TCGA dataset (Fig S5C).

Taken together, these analyses uncover *HAS2* as a stromal gene highly correlated with *NRG1* in luminal breast cancer patients.

### *NRG1* downregulation in CAFs downregulates *HAS2* and impairs their migration

We had observed that high-*NRG1* CAFs displayed higher proliferation rates and showed signatures of proliferation (Fig S4A and Table S3). We thus explored the potential contribution of *NRG1* expression to this phenotype in CAFs. To this end, we first knocked down *NRG1* in a high-*NRG1* CAF line by transient transfection. We selected two independent siRNA reagents with different degrees of downregulation to mimic heterogeneous downregulation levels (Fig 6A-B). Functional downregulation of *NRG1* was confirmed by using the CAF-siNRG1 CM on cancer cells. Both T47D and MCF7, showed a decrease in HER3 phosphorylation after incubation with CM from siNRG1 transfected CAFs, which paralleled the extent of *NRG1* downregulation in the CAFs (Sup Fig S6A). Similarly, migration of cancer cells induced by CAFs CM was also diminished upon *NRG1* downregulation in CAFs (Fig S6B). Having shown that the levels of *NRG1* downregulation in CAFs were sufficient to affect cancer cells, we next investigated the autocrine effect of NRG1 in CAFs. We observed that decreased NRG1 expression resulted also in a reduced proliferation rate in CAFs themselves (Fig 6C). Proliferation correlated with the efficiency of downregulation, however ectopic addition of NRG1 did not rescue that phenotype (Sup Fig S6C). This indicates that CAF-secreted NRG1 positively contributes to proliferation of cancer cells while it does not affect CAFs. Thus, we next investigated if the proliferative effect of NRG1 expression in CAFs was dependent on the binding of NRG1 to HER3 (Fig 3B). Strikingly, contrary to cancer cells, inhibition of NRG1 binding to HER3 by lumretuzumab treatment, did neither decrease proliferation of CAFs nor phosphorylation of AKT and ERK (Figs 6D and E). Altogether, these data demonstrate that the autocrine effect exerted by NRG1 on CAFs proliferation is independent of the canonical binding of secreted NRG1 to HER3. Further supporting this finding, expression levels of HER3 in CAFs were very low compared with expression levels in cancer cells and even lower in high-*NRG1* CAFs (Fig S6D). Of note, binding of NRG1 to HER4 receptor was not considered due to its undetectable expression in CAFs (Fig S6E).

**Figure 6.**
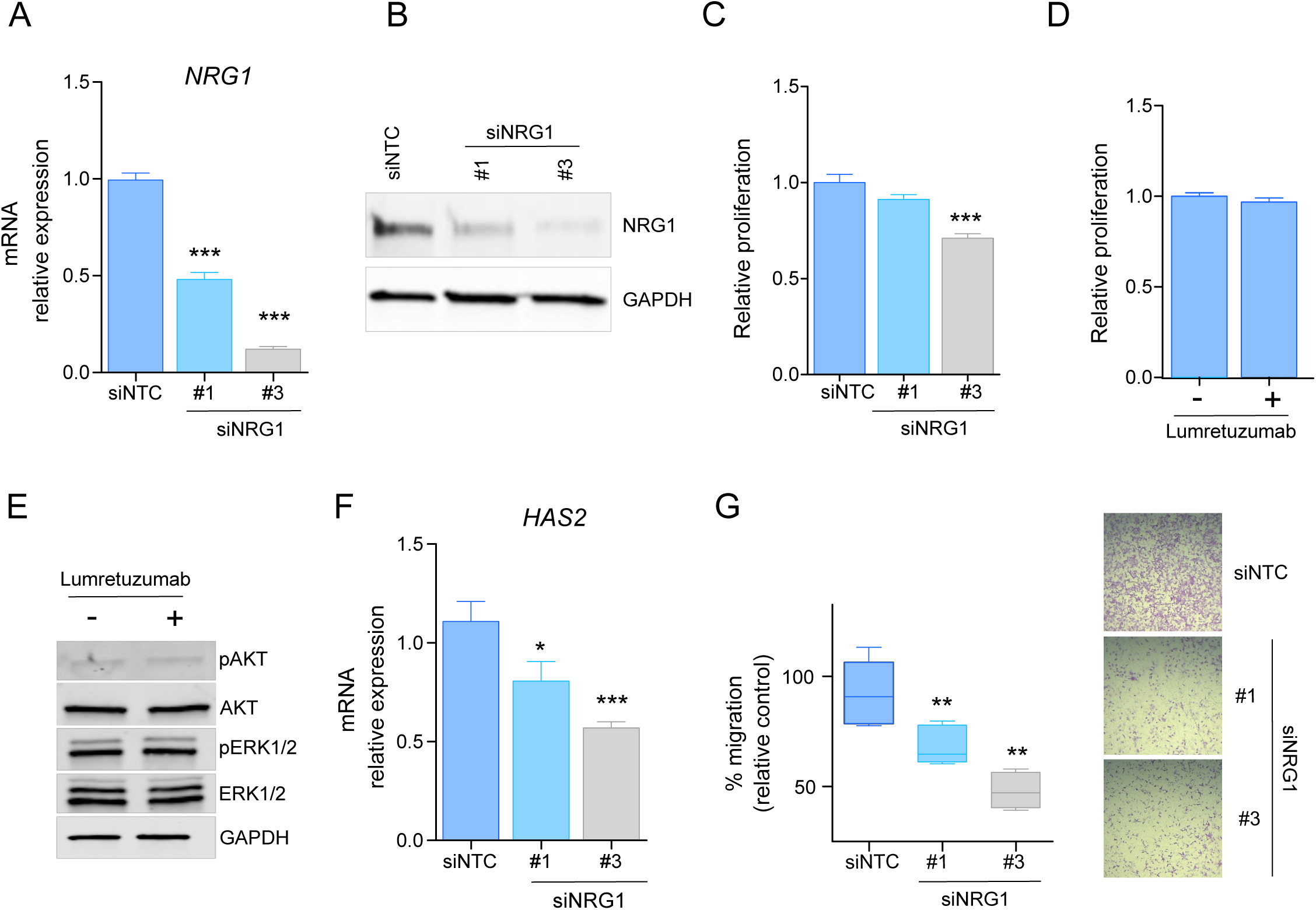
Functional implications of *NRG1* in CAFs. **A**, downregulation of *NRG1* at RNA level in CAF#3 (high-*NRG1*) using two independent siRNAs (siNRG1 #1, #3). Bar graphs represent average of 3 independent experiments and 3 technical replicates each. U-Mann Whitney test comparing with siRNA non-targeting control (siNTC) (\*\*\**P* < 0.01). **B**, western blot showing efficiency of downregulation of NRG1 at protein level in CAF#3 (high-*NRG1*). Two independent siRNA were chosen to obtain variable degrees of *NRG1* knockdown. **C**, relative cell growth of CAF#3 was measured 72h post re-seeding (5 days post-transfection) either with two different siRNAs targeting *NRG1* (siNRG1 #1, #3) or a non-targeting control siRNA (siNTC). U-Mann Whitney test was applied for statistical analysis (\*\*\**P* < 0.01). **D**, relative cell number of CAF#3 72h after treatment with 10μg/ml lumretuzumab (Lum). **E**, total protein and phosphoprotein levels for AKT and ERK1/2 after 24h of treatment with or without lumretuzumab (Lum) at 10μg/ml. GAPDH was used as loading control in CAF#3. **F**, *HAS2* mRNA levels in cells treated either with a non-targeting control siRNA (siNTC) or two different siRNAs targeting *NRG1* (siNRG1#1, #3). Bar graphs represent average of 3 independent experiments (n = 3 technical replicates). U-Mann Whitney test comparing with siRNA non-targeting control (siNTC) (\*\**P* < 0.01). **G**, migration of CAFs transfected either with a non-targeting control siRNA (siNTC) or two siRNAs targeting NRG1 (siNRG1#1, siNRG1#3) and normalized with seeding control. Whiskers in the box plot represent minimum and maximum values. U-Mann Whitney test comparing with siRNA non-targeting control (siNTC) (\*\**P* < 0.01). Crystal violet staining of transwell inserts representing migration after 8h of CAF#3 transfected either with a control siRNA (siNTC) or either of siRNAs targeting NRG1 (siNRG#1, siNRG1#3).

Transcriptomic profiling had revealed an enrichment of signatures related to migration in high-*NRG1* CAFs (Fig 5A). Additionally, *NRG1* correlated with *HAS2*, a known mediator of migration, in several CAF lines and patient stroma datasets (Figs 5B and C). We thus wondered if *NRG1* downregulation could affect not only proliferation but also *HAS2* levels and migratory capacity of CAFs. Remarkably, we observed a proportional decrease in *HAS2* mRNA transcript levels upon knockdown of *NRG1* (Fig 6F) which was associated with a marked decrease of the migration of CAFs (Fig 6G).

Collectively, this data suggests an autocrine, HER3-independent role of NRG1 in cancer-associated fibroblasts that modulates their proliferation and migration.

## Discussion

Overexpression of human epidermal growth factor receptor 3 (HER3) plays an important role in cancer development as well as acquired drug resistance in a wide variety of solid tumours (Karachaliou, Lazzari et al., 2017, Mishra, Patel et al., 2018). It has been associated with worse clinical outcome, and monoclonal antibodies such as lumretuzumab have been developed to neutralize its activity by blocking the binding of its ligand NRG1 (Mirschberger et al., 2013, Schneeweiss et al., 2018). Several preclinical and clinical studies have supported NRG1 as a predictive biomarker for anti-HER3 targeted therapies (Jacob et al., 2018, Liu, Liu et al., 2019, Ocana et al., 2016). NRG1-mediated autocrine signaling in cancer cells has been reported to underlie sensitivity to anti-HER2 therapies in certain ovarian and head and neck tumours (Sheng, Liu et al., 2010, Wilson et al., 2011). In our study, we show that the stromal compartment is the major contributor of NRG1 expression in breast cancer and it is not detected in luminal breast cancer cells (Chua et al., 2009). Our results suggest that in breast cancer, cells from the luminal subtype, depend on paracrine NRG1 to activate downstream pathways. Here, we used primary fibroblasts isolated from luminal breast cancer tissue and demonstrated that NRG1 secreted by CAFs is sufficient to activate the HER3 pathway in cancer cells. Activation of HER3 by CAF CM promotes phosphorylation of main downstream activators AKT and ERK, leading to proliferation and migration of cancer cells. The use of lumretuzumab, a humanized monoclonal antibody that selectively binds to the extracellular domain of HER3 thereby blocking binding of NRG1, is able to prevent that phenotype in a NRG1-dependent manner. Thus, we suggest that the utility of NRG1 as a predictive biomarker to anti-HER3 therapies in luminal breast cancer may be provided by the stromal compartment, while analysis of bulk tumour tissues may dilute its detection (Yoshihara, Shahmoradgoli et al., 2013).

It is widely accepted that CAFs are a heterogeneous population of mesenchymal cells defined by their diversity in functions, markers and origins. Several studies have compared gene expression in disease-free fibroblasts and CAFs derived from various tissues to obtain information on stromal pathways facilitating malignant phenotypes (Berdiel-Acer, Sanz-Pamplona et al., 2014, Saadi, Shannon et al., 2010, Sadlonova, Bowe et al., 2009). Other works have been oriented towards identifying specific lineages within CAFs based on their tumour promoting abilities to identify subpopulations (Costea, Hills et al., 2013, Morsing, Klitgaard et al., 2016, Patel, Vipparthi et al., 2018). Also, recent studies have described novel approaches for the study of biological function and targeting of CAFs (Su, Chen et al., 2018). Here, we have identified heterogeneous expression of *NRG1* in the stroma of luminal breast cancer tissue. Its higher expression categorizes and defines CAFs with an associated motile, fibrotic transcriptome and phenotype. Additionally, unsupervised clustering based on the proteomic profile of relevant signaling effectors such as ERK1/2, AKT, MEK, and c-JUN, classified CAF lines in the same high- and low-groups, reassuring the role of NRG1 in defining a different activation status.

The differential expression analysis conducted in this study revealed 102 genes upregulated in high-*NRG1* CAFs which were enriched in gene signatures related to a motile phenotype. We identified *ITGB2* and *EPHB1* as strongly correlating with *NRG1* expression in breast CAFs. Indeed, both *ITGB2* and *EPBH1* have been previously documented to play significant roles in polarization and cell migration (Karagiannis, Poutahidis et al., 2012).

Finally, we revealed *HAS2* (Hyaluronan Synthase 2) as a gene that is strongly correlated with *NRG1*, not only in primary CAFs but also in patient stroma datasets. HAS2 is responsible for the synthesis of hyaluronan (HA), a glycosaminoglycan with a demonstrated role in cancer initiation and progression and whose elevated accumulation in either the stroma or tumour parenchyma of many cancers is linked to tumour aggressiveness and poor outcome (Kim, Lee et al., 2019, Zhang, Tao et al., 2016). Functionally, its deficiency in mesenchymal stromal cells has been shown to be associated with an attenuation of CAF marker expression and poor migratory potential (Spaeth, Labaff et al., 2013). We suggest that in our system, correlation of *HAS2* and *NRG1* is consequence of a regulatory mechanism in which NRG1 expressed by CAFs regulates *HAS2* expression and modulates migratory potential of the fibroblasts by NRG1 non-canonical signaling (Mei & Nave, 2014, Mei & Xiong, 2008). Although additional molecular characterization will be necessary to further decipher the exact mechanism of this regulation in cancer, the strong correlation observed also in tumour stroma of patient samples clearly propose these two molecules as potential CAFs biomarkers. Thus, we consider that dual targeting of NRG1 and HAS2 may be an interesting treatment strategy for tumours with high expression of NRG1. Indeed, inhibition of *HAS2* upon treatment with its specific inhibitor 4-methylumbelliferone (4-MU), has proven successful in reducing tumour stroma in pancreatic ductal adenocarcinoma (PDAC) (Kudo, Suto et al., 2017, Yoshida, Kudo et al., 2018), which also shows high expression of NRG1 (Ogier et al., 2018). We hypothesize that, for breast cancer, on one side, treatment with anti-HER3 monoclonal antibodies would reduce the proliferation and migration of cancer cells by blocking stromal NRG1 binding, thus diminishing tumour aggressiveness. Concomitantly, the use of a specific HAS2 inhibitor would induce a reduction of the stroma content by decreasing HA synthesis by CAFs (Fig 7) further contibuting to tumour growth/aggressiveness wane. Ultimately, this tailored combination therapy represents a novel treatment approach from which NRG1-high breast cancer patients could benefit.

**Figure 7.**
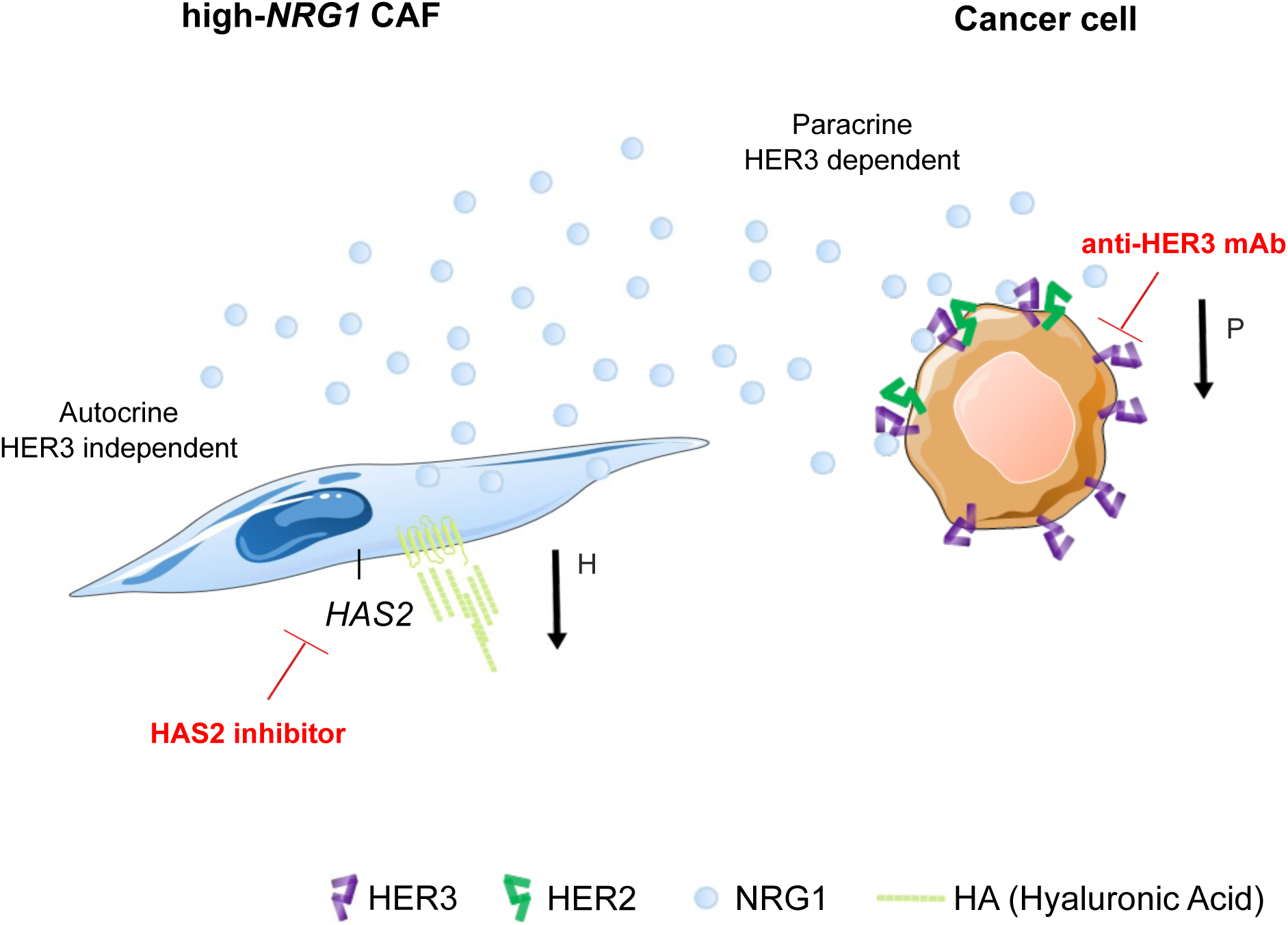
Model for dual targeting of high-*NRG1* stroma in luminal breast cancer. High-*NRG1* CAFs (blue) secrete high quantities of NRG1 (blue circles) promoting paracrine activation of HER3-receptor in cancer cells and thereby inducing proliferation and migration processes. Use of lumretuzumab blocks HER3 receptor in cancer cells and avoids binding of NRG1 thereby reducing proliferation and migration processes in the tumour cells. Suggested NRG1 non-canonical signaling induces proliferation and migration in CAFs, in a HER3-independent manner. CAFs are highly proliferative and migratory also due to increased expression of *HAS2* and secretion of HA. Additional use of a HAS2 inhibitor could help to reduce expression of *HAS2* and thus secretion of HA reducing migration of CAFs (/https://smart.servier.com/).

## Materials and Methods

### Cell culture

The human MCF7 (HTB-22) and T47D (HTB-133) luminal breast cancer cell lines were obtained from the American Type Culture Collection (ATCC, LGC Standards GmbH, Wesel, Germany) and maintained in Dulbecco’s Modified Eagle Medium: Nutrient Mixture F-12 (DMEM/F12), supplemented with 10% FCS, 50 units/ml penicillin and 50 µg/ml streptomycin sulfate (Invitrogen AG, Carlsbad, CA, USA) at 37°C with 5% CO2. The cell lines were authenticated by Multiplexion (Heidelberg, Germany) and negatively tested for mycoplasma contamination before and after completion of the study.

### Clinical samples, cell isolation and characterization

Tumour tissue was collected from patients (n = 6) undergoing surgery for breast carcinoma at the Department of Gynecology Martin-Luther-University Halle-Wittenberg in Halle (Saale), Germany, following ethical approval and written informed consent. To obtain CAFs, fresh breast cancer tissue was prepared by the pathologist under semi-sterile conditions and maintained refrigerated at 4°C in complete medium until its processing. Tumour tissue was mechanically minced into pieces (1-4 mm^3^) and centrifuged at 1600rpm for 10 min. After fat removal, pellet containing small pieces was resupended in DMEM/F12 (10% FCS, 1% P/S and 1% fungizone), filtered with a cell strainer (70µm) and plated in a 60mm culture dish. Outgrowth of cells was daily checked and medium renewed twice per week. After complete outgrowth in a 60cm^2^ dish, cells were passaged with 0.25% Trypsin-EDTA (Gibco, Life Technologies) and fibroblasts seeded into a new 100mm culture dish. After three cell passages, morphologically homogeneous cultures containing only fibroblasts were obtained and RNA and protein collected for further characterization. To obtain conditioned media, 2.5×10^5^ CAFs were seeded in 100mm culture dish. Once cells attached, media was replaced by DMEM/F12 (1% FCS, 1% P/S), incubated for 24h and collected for their further use.

### RNA isolation and analysis

Total RNA of primary CAFs and cancer cells was isolated with RNeasy Mini kit (Qiagen, Hilden, Germany) according to the manufacturer’s instructions. For mRNA, cDNA synthesis was carried out with the Revert Aid H Minus First Strand cDNA Synthesis Kit (Fermentas, Waltham, MA, USA). Quantitative RT-PCR (qRT-PCR) reactions for target genes were performed with the Applied Biosystems QuantStudio™ 3 & 5 Real-Time PCR System, using probes from the Universal Probe Library, UPL (Roche). The housekeeping genes *ACTB* and *GAPDH* were used for normalization of mRNA analysis. List of primers are provided in Table S7.

### Antibodies and immunoblotting

For Western blotting, cells were lysed in ice-cold RIPA lysis buffer (Thermo Fisher Scientific) containing protease inhibitor Complete Mini and phosphatase inhibitor PhosSTOP (Roche). Protein concentrations were determined by BCA Protein Assay Reagent Kit (Thermo Fisher Scientific) and proteins were denatured with 4xRoti Load (Carl Roth, Karlsruhe, Germany) at 95°C for 5 min. Depending on the size, proteins were separated by 12 or 15% SDS-PAGE, blotted onto a PVDF membrane Immobilon-FL (Merck Millipore, Darmstadt, Germany) and incubated with primary antibodies overnight at 4°C. List of antibodies is provided in Table S7. Secondary IRDye®680 or IRDye®800-conjugated antibodies (LI-COR, Lincoln, NE, USA) were used for band visualization. Membranes were scanned and analyzed with Odyssey scanner and Odyssey 2.1, respectively (LI-COR, Lincoln, NE, USA). For quantification, local background subtraction and GAPDH/Tubulin normalization were applied.

### Immunofluorescence

For detection of specific markers, 7.5×10^4^ fibroblasts were seeded in 6-well plate containing glass coverslips and cultured until 60-70% confluency. Cells were fixed with 4% paraformaldehyde for 10min at room temperature, permeabilized with 0.25% Triton X-100 for 10min and blocked with 3% BSA for 30min. Primary antibodies against α-smooth muscle actin (ab7817, 5µg/mL, Abcam) and FAP (ab53066, 1:150, Abcam), were incubated overnight at 4°C in a humidified chamber. 1h incubation with respective secondary antibodies containing DAPI (1:1000) was performed and images were acquired with Zeiss Cell Observer inverted microscope.

### Reverse Phase Protein Array (RPPA)

RPPA experiments were performed as previously described (Sahin, Lobke et al., 2007, Sonntag et al., 2014). Briefly, cell lysates from three biological replicates for each condition were spotted in nitrocellulose-coated glass slides (Oncyte Avid, Grace-Biolabs, Bend, OR, USA) in technical triplicates. All the primary antibodies used were previously validated through Western blots to test their specificity. Signal intensities of spots were quantified using GenePixPro 5.0 (Molecular Devices, Sunnyvale, CA, USA). Intensity values were log2 transformed and plotted using morpheus software (https://software.broadinstitute.org/morpheus/). List of antibodies used is provided in Table S7.

### Drug treatments

Lumretuzumab and pertuzumab were provided by Roche Diagnostics GmbH (Penzberg). Prior to addition of CAF-CM or human recombinant NRG-1β (4711, BioCat), cells were pre-treated with either lumretuzumab or pertuzumab (10μg/ml) for 30min in low serum media (1% FCS). For viability/proliferation assays, media was removed and CAF-CM or low serum media (with or without 50ng/ml NRG-1β) was added and incubated for 3 days. For short perturbation assays, incubation time was 5min prior to lysates collection.

### Viability and proliferation assays

3000 cancer cells were seeded in 96-well white plates and the effect of lumretuzumab, pertuzumab, human recombinant NRG-1β (4711, BioCat) or the different CM on cell viability was evaluated using CellTiter-Glo® luminescent assay (G7570, Promega). Prior to addition of CM or NRG-1β (50ng/ml), cells were pre-treated with 10μg/ml lumretuzumab or pertuzumab for 30min. Luminescence was determined after 3 days of culture using the GloMax® microplate reader (GM3000, Promega) and normalized with seeding control plate.

To determine proliferation rate, 1,000 fibroblasts or 3,000 cancer cells, were seeded in 96-well black plates and cell counting was measured by nuclei staining. Hoechst 1/10,000 (33342, Thermo Fisher), was added for 45 minutes prior to cell acquisition with MetaXpress microscope (Molecular Devices) at the desired time points.

### Migration assays

For scratch assay, 30,000 MCF7/T47D cells (in 10% FCS 1% P/S DMEM-F12 standard media) were seeded in 96-well black plates. Cells were allowed to grow until confluence, and starved O/N in 1% FCS media. The following day, cells were stained with Cell Tracker™ Green CMFDA (Invitrogen C2925, 10mM in DMSO) at a 1:5,000 dilution in starvation media for 30min at 37°C. Afterwards, media was replaced by fresh starvation media and incubated for additional 30-45min at 37°C to allow for the metabolization of the dye. By using a 96-well multichannel pipettor, a scratch was performed in each well. Wells were washed with PBS several times to remove any floating cell and 100μl/well of the corresponding conditions in 1% FCS were added. Images were acquired at the initial time point and after 21h with MetaXpress microscope (Molecular Devices). Images were analyzed using the MRI Wound Healing Tool in ImageJ (Schindelin, Arganda-Carreras et al., 2012), and the gap closure was determined by comparing the scratch surface areas at 0h and 21h post CM addition.

For migration of fibroblasts, 30,000 CAFs were seeded in the upper chamber of a Transwell plate (Corning® 3422, 8μm pore size) in media without FCS. Media containing 10% FCS was added to the bottom well. After 8h, cells were fixed with 4% PFA and stained with crystal violet (CV). Images were taken for the different conditions and elution of the crystal violet with acetic acid was used to quantify absorbance. Normalization was done by eluting the CV of the same number of cells seeded in an independent well.

### Transfections

Transfections with siRNA were performed using Lipofectamine RNAimax (Invitrogen) according to the manufacturer’s instructions. ON TARGETplus siRNAs were obtained from Dharmacon (Lafayette, CO, USA). For each gene, individual siRNAs were tested and two were selected for further experiments. ON TARGETplus nontargeting siRNA pool (Dharmacon) was used as control. For the siRNA screen, fibroblasts were seeded in 10cm dishes with their growth medium without antibiotics. Twenty-four hours later, cells were transfected with siRNA at a final concentration of 20nM. Twenty-four hours after transfection, fibroblasts were re-seeded for the different assays. Sequences of siRNA used are in Table S7.

### Analysis of expression datasets (LCM)

For the analysis of the GEO datasets with the accession numbers: GSE10797 (Casey et al., 2009), GSE14548 (Ma et al., 2009), GSE35019 (Vargas et al., 2012) and GSE83591 (Liu et al., 2017), normalized data were downloaded from GEO.

### NGS data preprocessing and normalization

Whole genome RNA sequencing was performed at the Genomics Core Facility of German Cancer Research Center (DKFZ - https://www.dkfz.de/gpcf) using the Illumina HiSeq 2000 platform. After a thorough quality control check NGS data was assembled, aligned & annotated to the human genome hg38 and normalized using the TPM (Transcript Per kilobase Million reads) method.

### Identification of highly variable genes

In order to identify the most variable genes in the CAF lines, the deviation of gene expression levels from a fitted regression line estimated on coefficient of variation of control feature expressions were obtained using generalized linear models (Nelder, 1989, Trevor J. Hastie, 2017). To control the effects of outlier in the data, a winsorization procedure was used on the expression matrix. 517 highly variable genes were identified via Χ^2^ test at *FDR* ≤ 0.001.

### Differential expression analyses

Differential expression analyses within CAF samples were performed using DESeq2 method (Anders & Huber, 2010, Love, Huber et al., 2014) utilizing factor scaling normalization via effective library size estimation and a negative binomial test. Conditions are defined by a design matrix, and corrections for multiple testing was done using Benjamini & Hochberg’s method (Y.Hochberg, 1995). Significant differentially expressed genes were considered those with *FDR*BH ≤ 0.05. (adj *P* value < 0.05, absolute logFC > 0.5).

### Functional analyses: Gene ontology

We used the BioInfoMiner online platform to investigate which biological process Gene Ontology (GO) terms (Ashburner, Ball et al., 2000, Harris, Clark et al., 2004) were enriched in the list of differentially expressed or highly variable genes. BioInfoMiner exploits biological hierarchical vocabularies by mapping the genes to a genomic network created from semantic data. It prioritizes them based on the topological properties of the network after minimising the impact of semantic noise (bias) through different types of statistical correction. It detects and ranks significantly altered processes and the driver genes involved. The BioInfoMiner platform is available online at the website https://bioinfominer.com.

### Transcription Factor enrichment analysis

The transcription factor enrichment analysis was performed using the ChEA_2016 database (Lachmann, Xu et al., 2010) with the Enrichr gene list enrichment analysis tool (Chen, Tan et al., 2013, Kuleshov, Jones et al., 2016) and selecting the significantly enriched subset of transcription factors that included NRG1 in their targets.

### Statistical analyses

Statistical analyses and graphical representation were performed using GraphPad Prism version 6.00 for Windows.

## Author Contributions

MBA designed and performed most of the experimental work in the study. AM and ZH carried out immunofluorescence and migration experiments, respectively. SB performed and analysed RPPA experiments. SB and CB assisted with experimental work. MV, DB and CT prepared human tumour samples and conducted pathological analysis. KA, IB and AC performed the bioinformatics analysis of RNA-seq data. MH provided materials and insights to the study. EE performed data interpretation and revised the manuscript. MBA and SW designed the study and wrote the manuscript.

## Acknowledgements

We thank N.Erdem and C.Koerner for valuable discussions, the Department of Gynecology, Martin-Luther-University Halle Wittenberg, Halle (Saale), Germany for providing tumour tissue and clinical data, as well as the Breast Cancer Now Tissue Bank (https://breastcancernow.org/) for providing primary fibroblasts. E.Reinz and S.Kemmer for contributing to Fig S1C. D.Heiss and S.Karolus provided excellent technical assisstance. The Genomics & Proteomics as well as the Light Microscopy Core Facilities at German Cancer Research Center performed excellent services. This work was supported by the German Federal Ministry of Education and Research (e:Med FKZ: 031A429).

## Supplementary figures

**Figure S1. *HER3* expression in luminal subtypes and paracrine activation**

**A**, expression of *HER3* in breast cancer subtypes extracted from METABRIC and TCGA datasets. Boxes indicate mean +/- quartiles and minimum and maximum values represented by bars. Statistically significant comparisons with luminal subtypes are depicted. ANOVA multiple comparison test (\**P* < 0.05, \*\**P* < 0.01,\*\*\*\**P* < 0.0001).

**B**, mRNA expression of HER receptors in breast cancer cell lines representing different subtypes; T47D, MCF7 (luminal A), BT474 (luminal B), MDA-MB-468 (triple negative) breast cancer cell lines. HER1-3 expression levels were normalized to the respective levels in MCF10A, a non-transformed breast epithelial cell line. HER4 expression levels were normalized to BT474.

**C**, protein intensities of HER3, AKT, ERK1/2 and their respective phosphorylation states in T47D and MCF7 cancer cell lines along time and with lumretuzumab (red), pertuzumab (blue) or solvent (black), obtained by reverse phase protein array (RPPA). Different phosphorylation dynamics are observed for MCF7 and T47D under treatment, mainly for ERK1/2.

**Figure S2. Characterization of carcinoma-associated fibrobasts (CAFs)**

**A**, immunofluorescence of common activation markers αSMA (red, upper panel) and FAP (red, lower panel) in T47D and MCF7 cancer cells used as negative control. Nuclei counterstaining with DAPI (blue). Representative images are shown.

**B**, mRNA values of α-smooth muscle actin (*ACTA2*) and fibroblast activation protein (*FAP*) in all fibroblasts relative to CAF#4. Bars represent mean +/- s.e.m of two independent experiments (n = 3 technical replicates each).

**C**, representative western blot of NRG1 protein in all CAFs and in two luminal breast cancer cell lines. Tubulin was used as a loading control.

**Figure S3. Activation of HER3 in cancer cells by secreted NRG1 is CAF-dependent**

**A**, heatmap representing relative phosphorylation values of HER3, AKT and ERK1/2 in T47D and MCF7 quantified by RPPA. Low-serum condition media (1% FCS) was used as negative control (Ctrl) and ectopic NRG-1β (50ng/ml) as a positive control. Controls correspond to the proteomic profile of T47D and MCF7 exposed to CM from CAFs with or without lumretuzumab in Fig 3A. **B**, relative proliferation of MCF7 and T47D cancer cells 72h after culture with conditioned media (CM) from different CAFs. Values are relative to the control (1% FCS, DMEM-F12 media). Graph bar represents mean +/- s.e.m of 3 independent replicates (n = 3 technical replicates). ANOVA multiple comparison test was performed among all CM and the control (\**P* < 0.05, \*\**P* < 0.01,\*\*\**P* < 0.0001).

**Figure S4. Heterogeneous expression of *NRG1* defines two clusters of CAFs**

**A**, number of CAFs quantified by nuclei staining (Hoechst) along 5 consecutive days in 10% FCS media. Each dot represents average of 2 independent replicates (n = 6 technical replicates each). Blue lines/dots corresponding to high-*NRG1* fibroblasts. Low-*NRG1* fibroblasts represented in grey.

**B**, ranking of fibroblasts based on *NRG1* expression (TPM - transcript per milion) RNA seq data. CAFs were designated as low-*NRG1* (CAF#4, CAF#1 and CAF#5, grey) and high-*NRG1* (CAF#6, CAF#2 and CAF#3, blue) based on mean *NRG1* expression.

**C**, graph bar representing adjusted *P* value and number of transcription factors in high-*NRG1* CAFs overlaping with the ChEA database.

**Figure S5. Genes correlating with *NRG1* expression**

**A**, correlation matrix of *NRG1* and associated genes. Matrix represents those genes with an absolute Pearson correlation > 0.8.

**B**, expression correlation of *NRG1* and markers *CD36, ACTA2,S100A4* and *WNT5A* in isolated CAFs based on TPM values. Spearman correlation *r* and *P* value are indicated.

**C**, correlation of *NRG1* and candidate genes in TCGA dataset samples. Only samples with tumour purity < 0.5 are represented. Spearman correlation r and *P* value are indicated per each gene/dataset.

**Figure S6. Downregulation of *NRG1* in CAFs**

**A**, protein levels of HER3, AKT, ERK1/2 and their phosphorylation status upon 5 min addition of CM from CAF#3 transfected with either siNRG1#1, siNRG1#3 or siRNA non-targeting control (siNTC). Western blot representative of 2 independent experiments.

**B**, cancer cells and percentage of closure after 21h in CM from CAF#3 transfected with either siNRG1#1, siNRG1#3 or siNTC. Student’s t-test comparing individual siRNA NRG1 with siNTC (\*\**P* < 0.01; \*\*\**P* < 0.001).

**C**, graph bar representing proliferation of CAF#3 line upon transfection with siRNA non-targeting control (siNTC), siNRG1#1 or siNRG1#3 without or with ectopic NRG1 (doted bars). No significance rescue of proliferation was obtained when addition of ectopic NRG-1β in any of the conditions.

**D**, mRNA expression levels of HER receptors in CAF lines normalized to expression of respective receptors in the T47D luminal cancer cell line.

**E**, dot plot representing the expression of HER receptors in low-*NRG1* (grey circles) and high-*NRG1* (blue circles) CAFs obtained by RNA-sequencing.

**Table S1. Pathological characteristics of breast cancer specimens**

Selected histopathological details of the tumour samples. Staging of the tumours was done according to the TNM classification (Sobin, 2011). Tumours were characterised based on positive (pos) or negative (neg) expression of biomarkers like estrogen receptor (ER), progesterone receptor (PgR) and HER2. ER expression was considered positive when > 1% of cells were stained. Percentage of cells staining positive for Ki67 is listed.

**Table S2. List of most variable genes (MVG)**

A total of 517 genes were identified as statistically variable (*FDR* ≤ 0.001) among 6 CAFs under study.

**Table S3. Biological processes represented by MVG**

Gene Ontology of the most variable genes (MVG) reveals the biological processes in which those fibroblasts are contrasting. Processes related with locomotion, migration and cell motility are highly represented.

**Table S4. Transcription factors enriched in high-*NRG1* CAFs**

Enriched transcription factors and their adjusted p value in high-*NRG1* CAFs, obtained from the ChEA_2016 database.

**Table S5. List of differential expression genes between high and low-NRG1 CAFs**

Genes with and adj *P* value < 0.05 and absolute logFC > 0.5 are listed. Total of 102 genes were upregulated in the high-*NRG1* group and 151 genes upregulated in low-*NRG1*.

**Table S6. Correlation matrix of NRG1-associated genes**

Pearson correlation among the 517 most variable genes.

**Table S7. Materials and methods**

**A**, description of antibodies used for RPPA and Western blot.

**B**, sequences and probes for RNA expression analysis.

**C**, siRNA target sequence for *NRG1* and *HAS2*. 2 independent siRNA were selected for each gene. A pool of 2 sequences of non-targeting siRNA used as a control.

